## Supplementary File for "Bioengineered niches that recreate physiological extracellular matrix organisation to support long-term haematopoietic stem cells"

### Supplementary figures

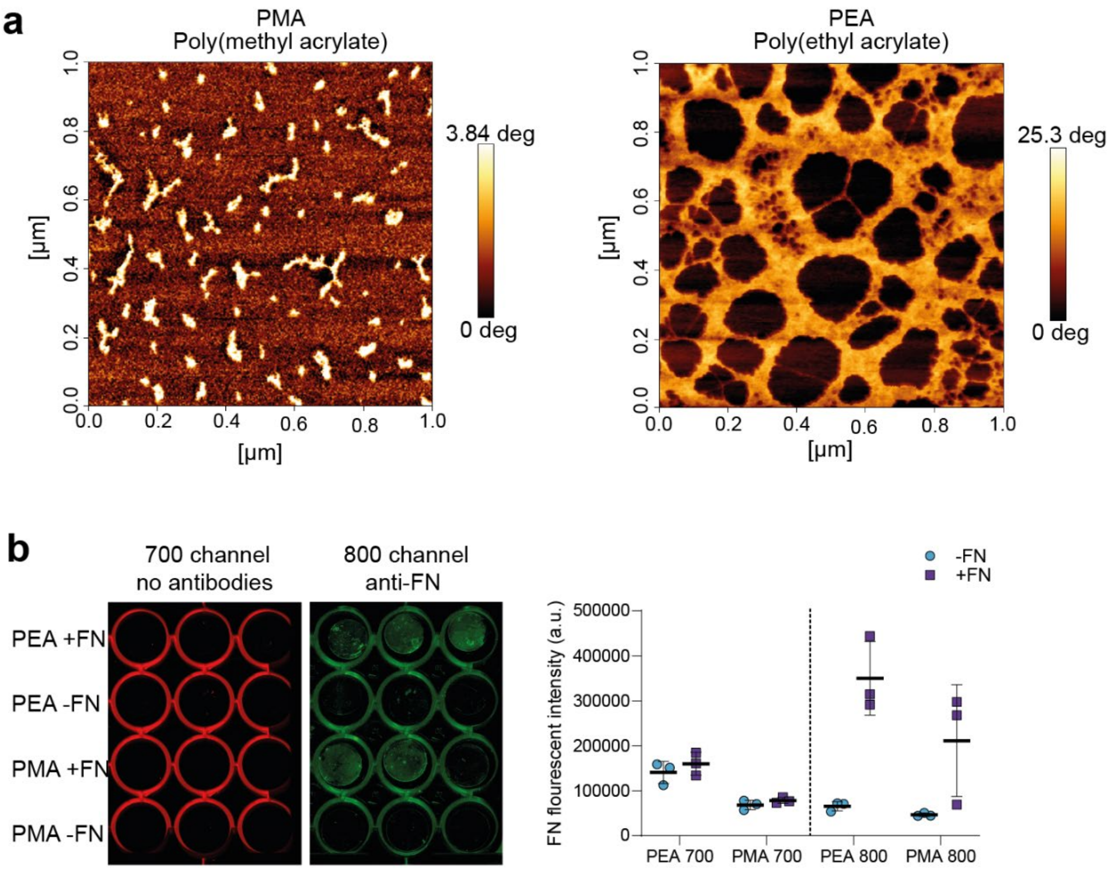


***Supplementary Figure 1 | Fibronectin (FN) quantification***. ***a***, *phase images for AFM corresponding to Figure 2b.* ***b***, *immunofluorescence analysis of PEA/PMA background fluorescence and FN quantification using in-cell western, demonstrates minimal background fluorescence observed in 700 channel containing no secondary antibodies, and 800 channel where there is no primary antibody target.*

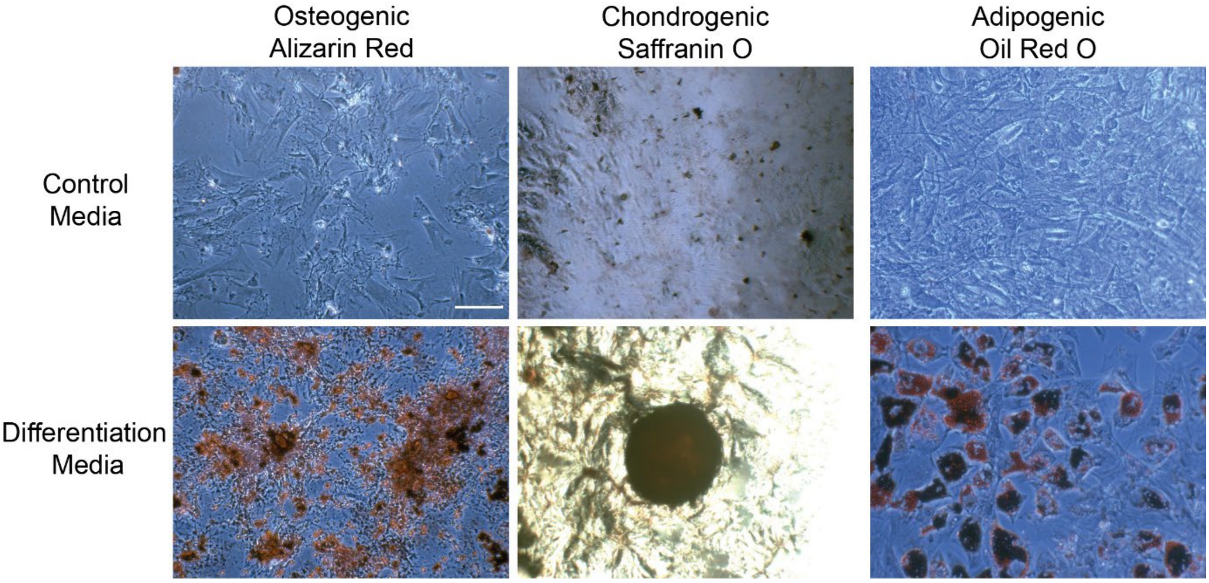


***Supplementary Figure 2 | Trilineage differentiation potential of PerSCs.*** *PerSCs were isolated and cultured ≤passage 5, then cultured in either control media or medias for osteogenic, chondrogenic and adipogenic differentiation of stromal stem cells. Histological stains were then used to assess differentiation after 4 wk culture. Alazarin red stains calcium deposits in osteogenic cultures. Safranin O is used to identify chondrocytes. Oil red O stains lipids in adipogenic cultures. Positive detection of all stains in differentiation medias demonstrates trilineage potential of PerSCs. Scale bar is 100 µm.*

**a**

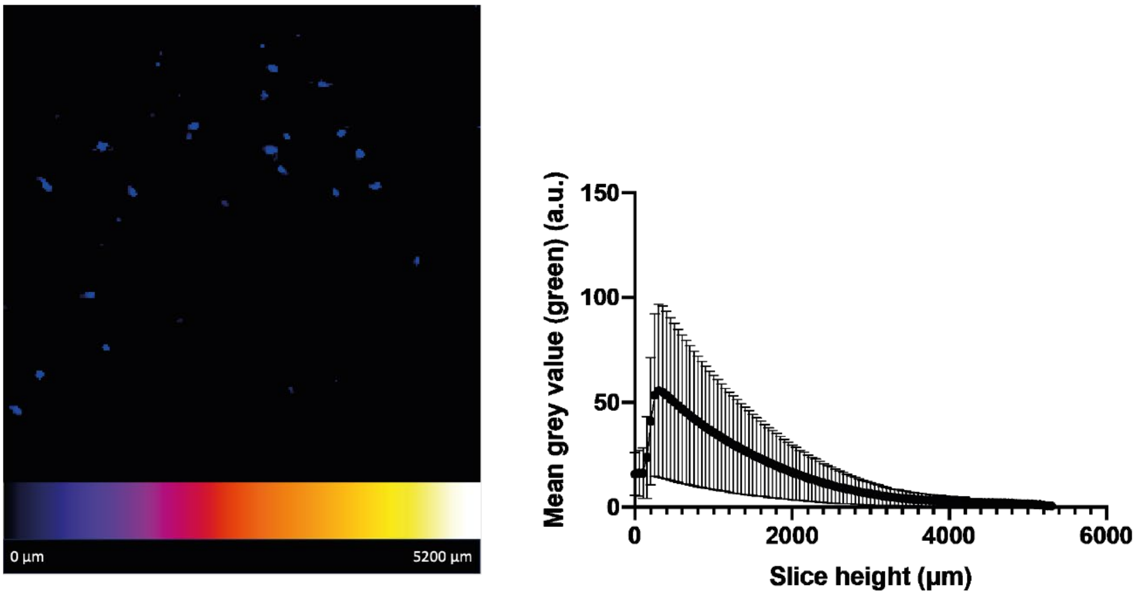


**b**

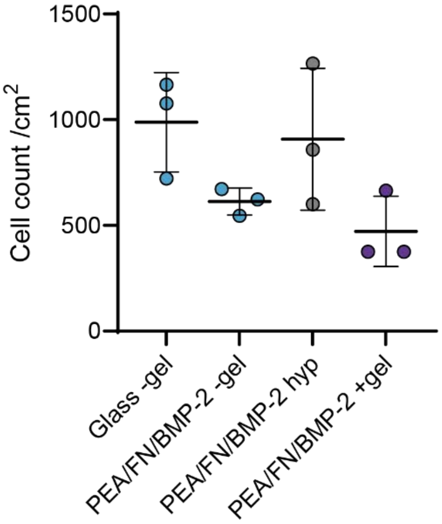


***Supplementary Figure 3 | PerSC quantification. a,*** *PerSC migration into collagen gels was assessed via Hoescht staining of PerSCs in the PEA +gel niche model. Z-projections show minimal migration into the gel, with most cells remaining in the bottom 2000 µm of the well.* ***b,*** *Cell number quantification, via Hoescht staining. Statistics by one-way ANOVA followed by Bonferroni multiple comparison test show no significant differences. Graph shows mean ± SD, n= 3 material replicates.*

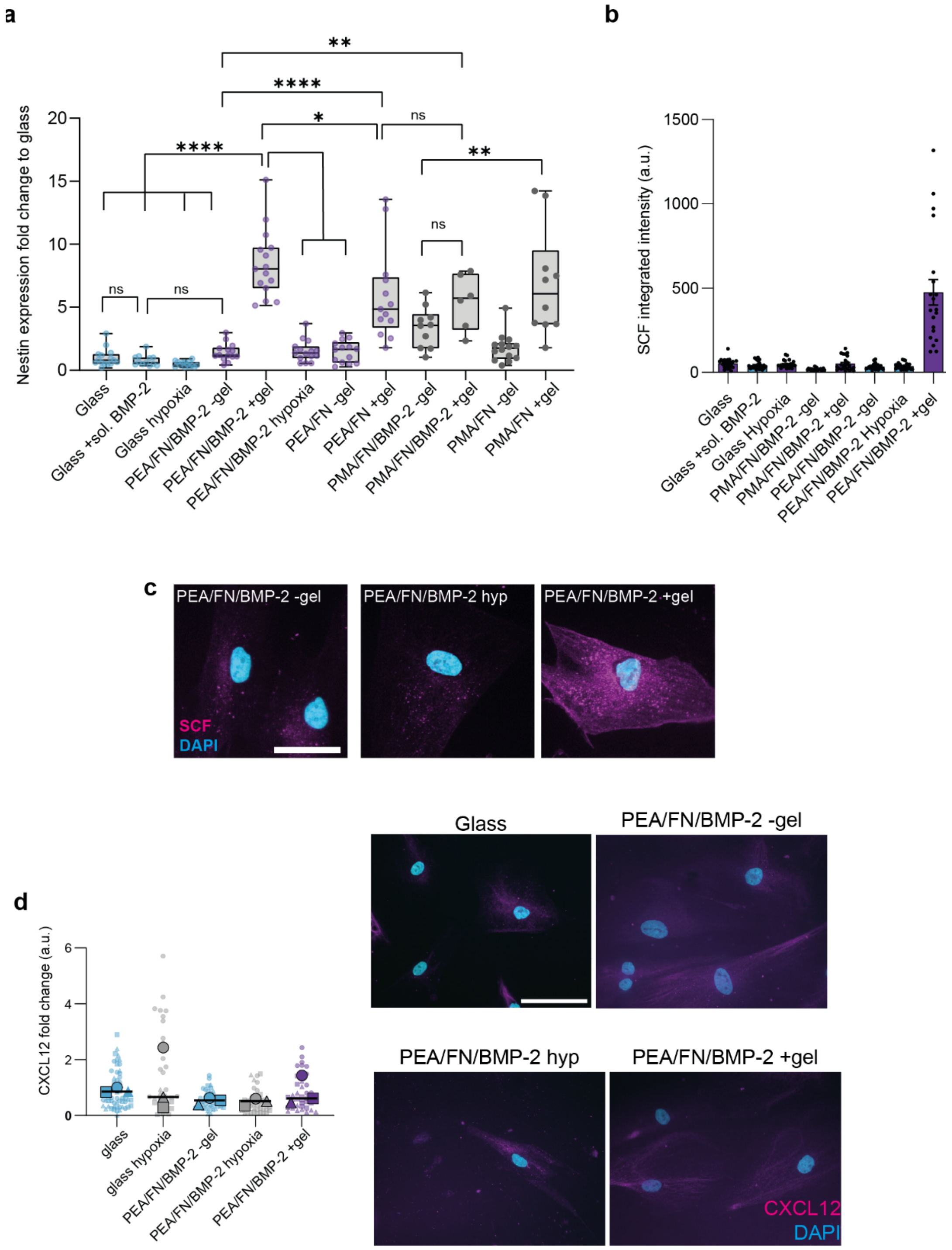


***Supplementary Figure 4 | Perivascular stromal cell niche phenotype. a****, Nestin expression measured by immunofluorescence at day 14 of culture. Nestin expression significantly increased in perivascular stromal cells on the PEA/FN/BMP-2 system only in the presence of low stiffness gels (+gel niche). PMA is used as a control polymer and shows some increase with gel only by day 14. Glass with soluble BMP-2 (50 ng/ml) added to the media is used as a further control and shows the addition of soluble, not matrix bound, BMP-2 does not induce nestin expression. Graph shows mean integrated intensity as fold change to glass control ± SEM, ****= p<0.001, **= p<0.005, determined by one-way ANOVA followed by Bonferroni multiple comparison test. N = 4 material replicates. Each point represents one field of view, normalised to cell number. We note a small trend of increase as we switch BMP-2 from soluble (glass +sol. BMP-2) to solid-phase (PEA-FN-BMP-2 -gel).* ***b, c,*** *SCF and* ***d****, CXCL12 production in niche models. Cells were cultured in niche systems for 14 days with brefeldin A treatment (5 $\mu$g/ml in culture media) added for the final 24 h to inhibit intracellular protein transport.*

*Quantification of immunofluorescence staining with b, anti-SCF shows significant increase only in the PEA/FN/BMP-2 +gel condition. Graph shows mean ± SEM, ****= p<0.001, determined by one-way ANOVA followed by Tukey's multiple comparison test. Each point represents integrated SCF intensity of 1ximage field normalised to cell number, from 4 material replicates.* ***c****, shows representative immunofluorescence images for SCF, corresponding the B and Figure 3e. For* ***d****, all conditions expressed detectable CXCL12, but with no significant differences. Graph shows mean integrated intensity CXCL12 as fold change to glass control (n=3 material replicates, for 3 independent experiments with different donor cells), each point represents 1x image field normalised to cell number with background correction. Representative images from 1 donor shown, scale bar is 100 µm, magenta = CXCL12, cyan = DAPI. Non-significant, determined by one-way ANOVA followed by Bonferroni multiple comparison test.*

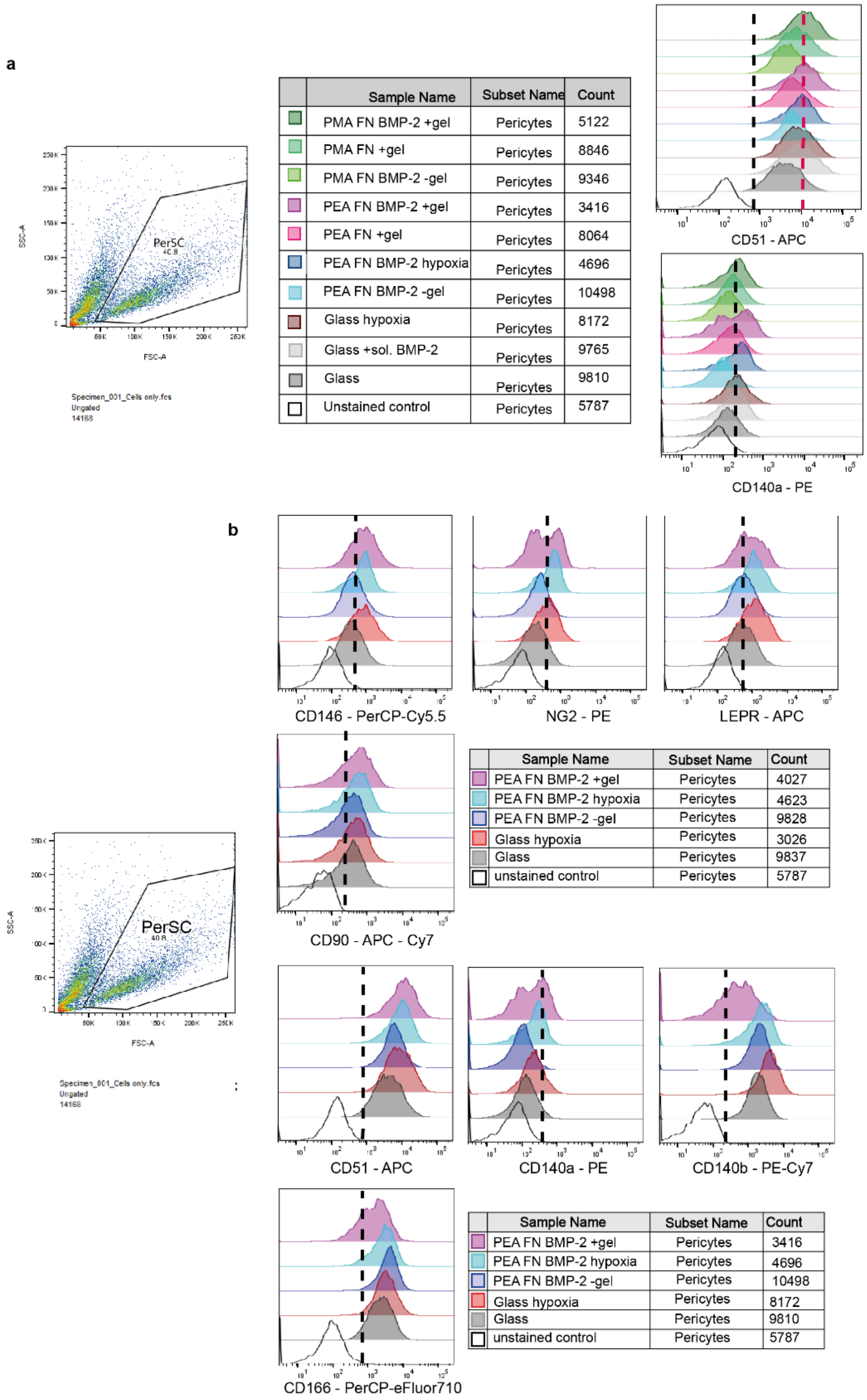


| Sample Name | Subset Name | Count |
|---|---|---|
| PMA FN BMP-2 +gel | Pericytes | 5122 |
| PMA FN +gel | Pericytes | 8846 |
| PMA FN BMP-2 -gel | Pericytes | 9346 |
| PEA FN BMP-2 +gel | Pericytes | 3416 |
| PEA FN +gel | Pericytes | 8064 |
| PEA FN BMP-2 hypoxia | Pericytes | 4696 |
| PEA FN BMP-2 -gel | Pericytes | 10498 |
| Glass hypoxia | Pericytes | 8172 |
| Glass +sol. BMP-2 | Pericytes | 9765 |
| Glass | Pericytes | 9810 |
| Unstained control | Pericytes | 5787 |

| Sample Name | Subset Name | Count |
|---|---|---|
| PEA FN BMP-2 +gel | Pericytes | 4027 |
| PEA FN BMP-2 hypoxia | Pericytes | 4623 |
| PEA FN BMP-2 -gel | Pericytes | 9828 |
| Glass hypoxia | Pericytes | 3026 |
| Glass | Pericytes | 9837 |
| unstained control | Pericytes | 5787 |

| Sample Name | Subset Name | Count |
|---|---|---|
| PEA FN BMP-2 +gel | Pericytes | 3416 |
| PEA FN BMP-2 hypoxia | Pericytes | 4696 |
| PEA FN BMP-2 -gel | Pericytes | 10498 |
| Glass hypoxia | Pericytes | 8172 |
| Glass | Pericytes | 9810 |
| unstained control | Pericytes | 5787 |

***Supplementary Figure 5 | Histograms of cell surface marker phenotyping**. Corresponding to Figure 3f. **a**, CD51 and CD140a – pseudo-markers for nestin expression[3] – are increased and show a double population (respectively) in nestin$^{high}$ PerSCs in PEA/FN/BMP-2 +gel niches . **b**, shows representative histograms from one representative biological replicate for the heatmap in Figure 3f.*

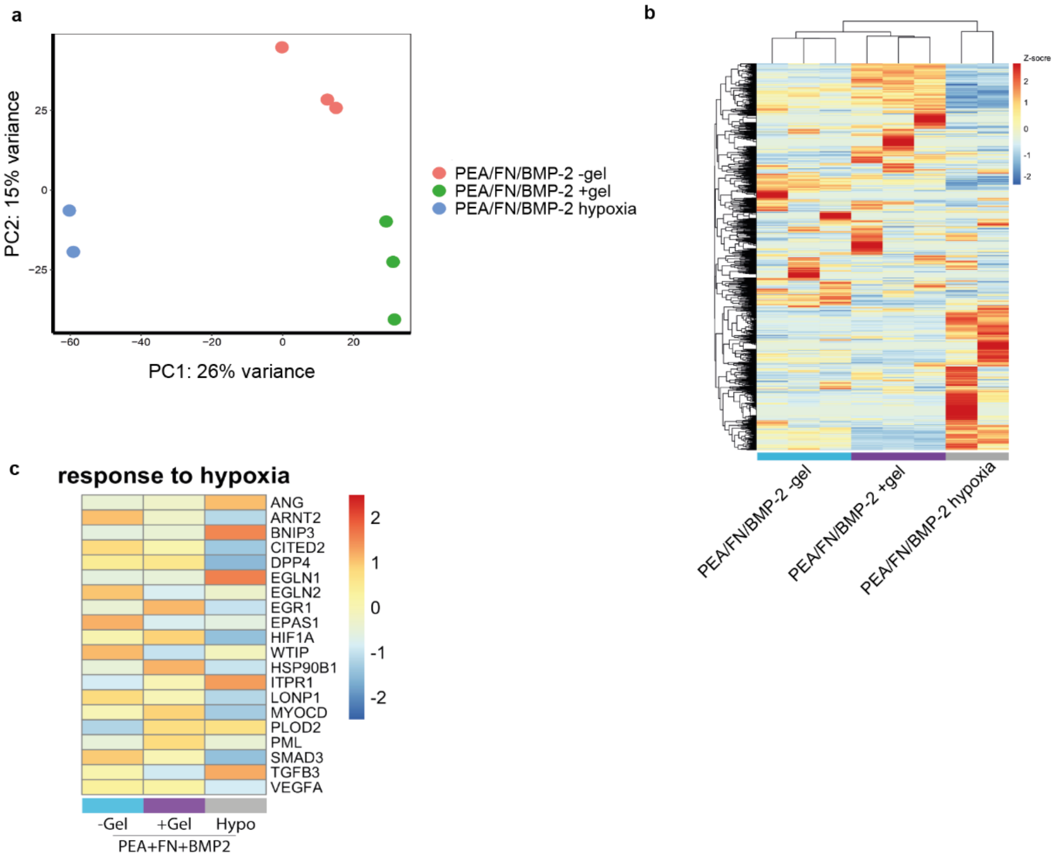


***Supplementary Figure 6 | RNA-seq analysis of PerSC niche phenotype at day 7. a,*** *Principal component analysis (PCA) shows transcripts from conditions PEA/FN/BMP-2 -gel/+gel/hypoxia samples cluster independently.* ***b****, Heatmap of z-scored changes in transcript levels.* ***c****, z-scored changes in transcript levels of genes involved in GO enrichment pathway 'response to hypoxia'. Genes with FDR < = 0.05 were considered as differentially enriched. N = 3 material replicates, for hypoxia n=2 material replicates as one replicate failed RNA quality control.*

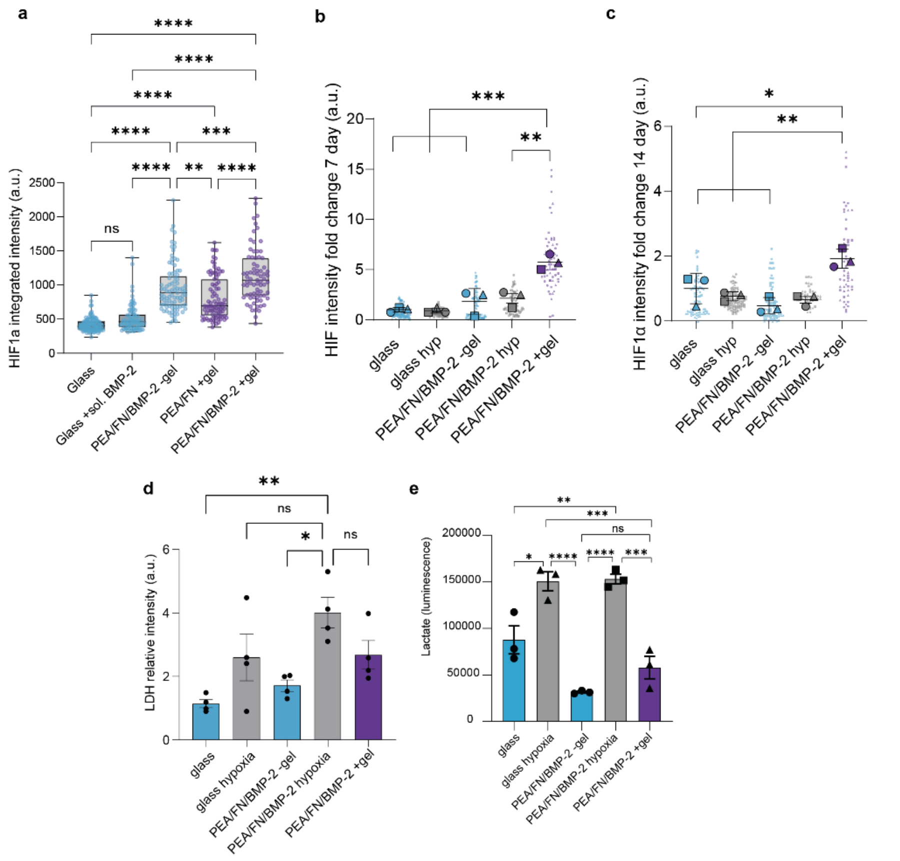


***Supplementary Figure 7| HIF1a co-localisation and downstream lactate regulation. a,*** *HIF1a co-localisation to nucleus, comparison to additional controls PEA/FN +gel and glass +soluble BMP-2, shows PEA/FN/BMP-2 +gel leads to significantly higher levels of nuclear HIF1a at 3 days. Data from one experimental repeat that is included in Figure 5a, with 3 material replicates, statistics by one-way ANOVA with Bonferroni multiple comparisons test. We note a significant increase as we switch BMP-2 from soluble (glass +sol. BMP-2) to solid-phase (PEA-FN-BMP-2 -gel).* ***b*** *and* ***c*** *Nuclear HIF1α levels were compared at day 7 and 14 by immunofluorescence microscopy. HIF1α levels increased in cells in all models, and significantly increased in +gel niches. Each point represents 1 nuclei measurement with shape corresponding to mean shape for 3 material replicates. Graph shows means ± SEM.* ***d*** *and* ***e*** *Lactate dehydrogenase (LDH) and lactate levels are not raised with PEA/FN/BMP-2 +gel.* ***d,*** *in-cell western analysis of LDH levels in cells detected by immunofluorescence at day 7. PerSCs in PEA/FN/BMP-2 with hypoxia had significantly increased levels of the enzyme.  Graph shows fluorescent intensity of LDH normalised to cell number (CellTag800) ± SEM, *= p<0.05, **= p<0.005, determined by one-way ANOVA followed by Bonferroni multiple comparison test. N = 4 material replicates.* ***d,*** *Lactate levels in cell supernatant at day 14 measured by luminescence. Glass and PEA/FN/BMP-2 hypoxia samples had significantly raised levels. Graph shows n= 4 material replicates, ± SEM, **= p<0.005, ***= p<0.001, ****= p<0.001 determined by one-way ANOVA followed by Bonferroni multiple comparison test.*

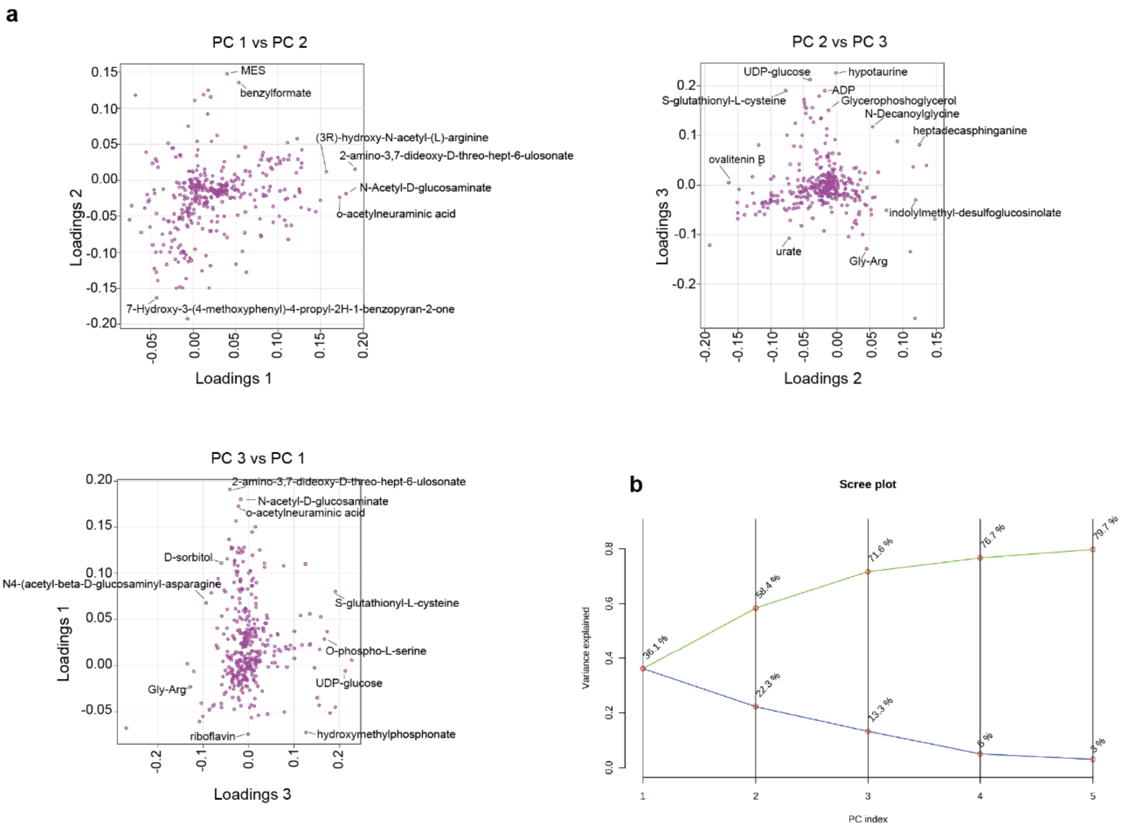


***Supplementary Figure 8 | Metabolic variables in principle components.*** ***a****, Loading plots for PCA of all metabolites with most variable metabolites labelled.* ***b****, Scree plot shows variance for each principal component.*

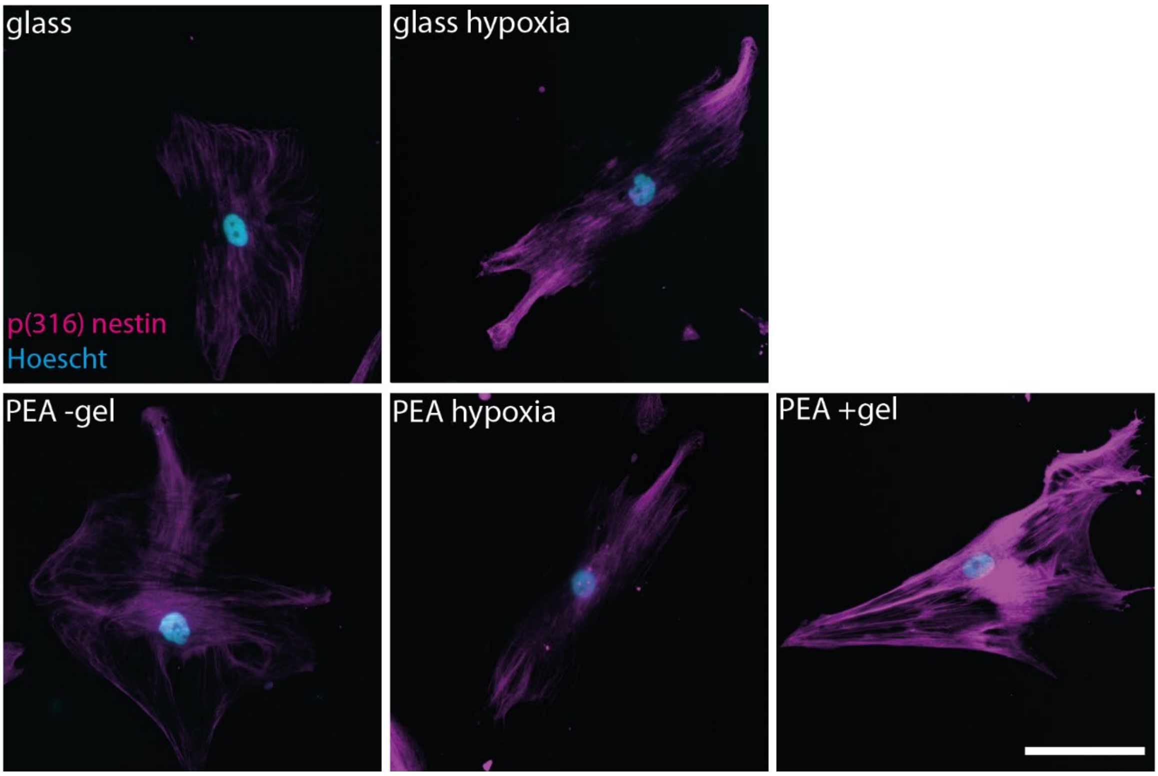


***Supplementary Figure 9 | Immunofluorescent detection of nestin phosphorylation at Th(316).*** *Representative images for Figure 6a. Suggests some nestin is phosphorylated and therefore soluble. Scale bar = 100 µm, magenta = p(316) nestin, cyan = Hoescht, n= 4 material replicates.*

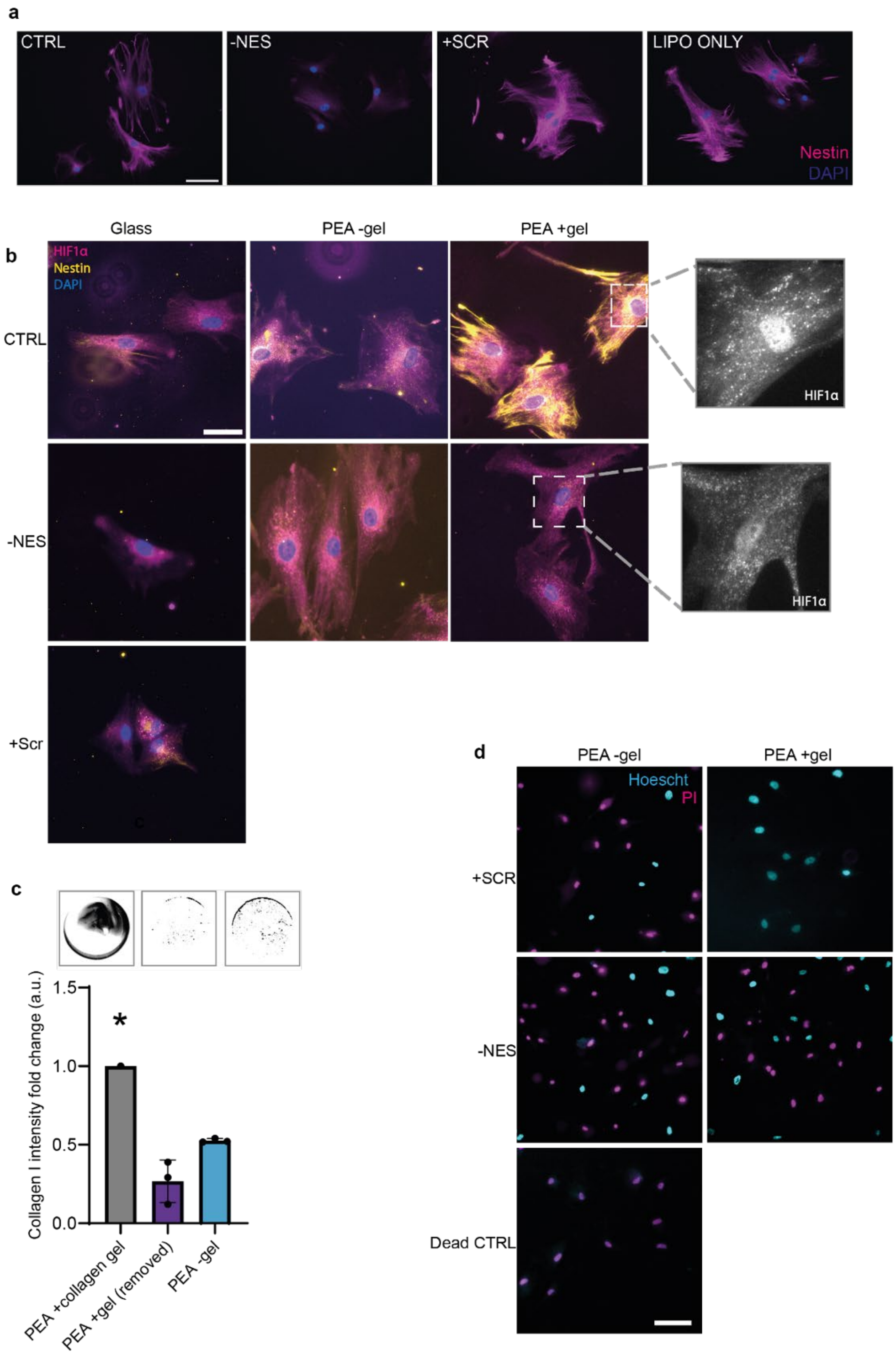


**Supplementary Figure 10 | Nestin knock-down by siRNA**. **a**, representative images for Figure 6b, demonstrating siRNA knock-down of nestin in PerSCs. Scale bar = 100 µm, magenta = nestin, blue = DAPI. Scr = scrambled control; lipo only = lipofectamine only control. **b**, Nestin silencing leads to loss of HIF1α nuclear co-localisation in PerSCs cultured in PEA/FN/BMP-2 +gel niches. Representative images for Figure 6c. Scale bar = 100 µm, magenta = HIF1α, yellow = nestin, blue = DAPI. **c**, Collagen I staining on fixed (but not permeabilised) PEA +PerSC coverslips after 7 days culture. Fluorescence intensity demonstrates mechanical gel removal for cytotoxicity experiments is complete and does not leave residual collagen. PEA +collagen gel (without removal) was used as a positive control. **d**, siRNA knock-down of nestin in PerSCs and susceptibility to oxidative stress was assessed. Representative images of 6d. All conditions were +$H_2O_2$, dead CTRL +10 mM $H_2O_2$. N= 4 material replicates, scale bar = 100 µm. Cyan = Hoescht (live and dead nuclei stain), magenta = PI (dead nuclei stain).

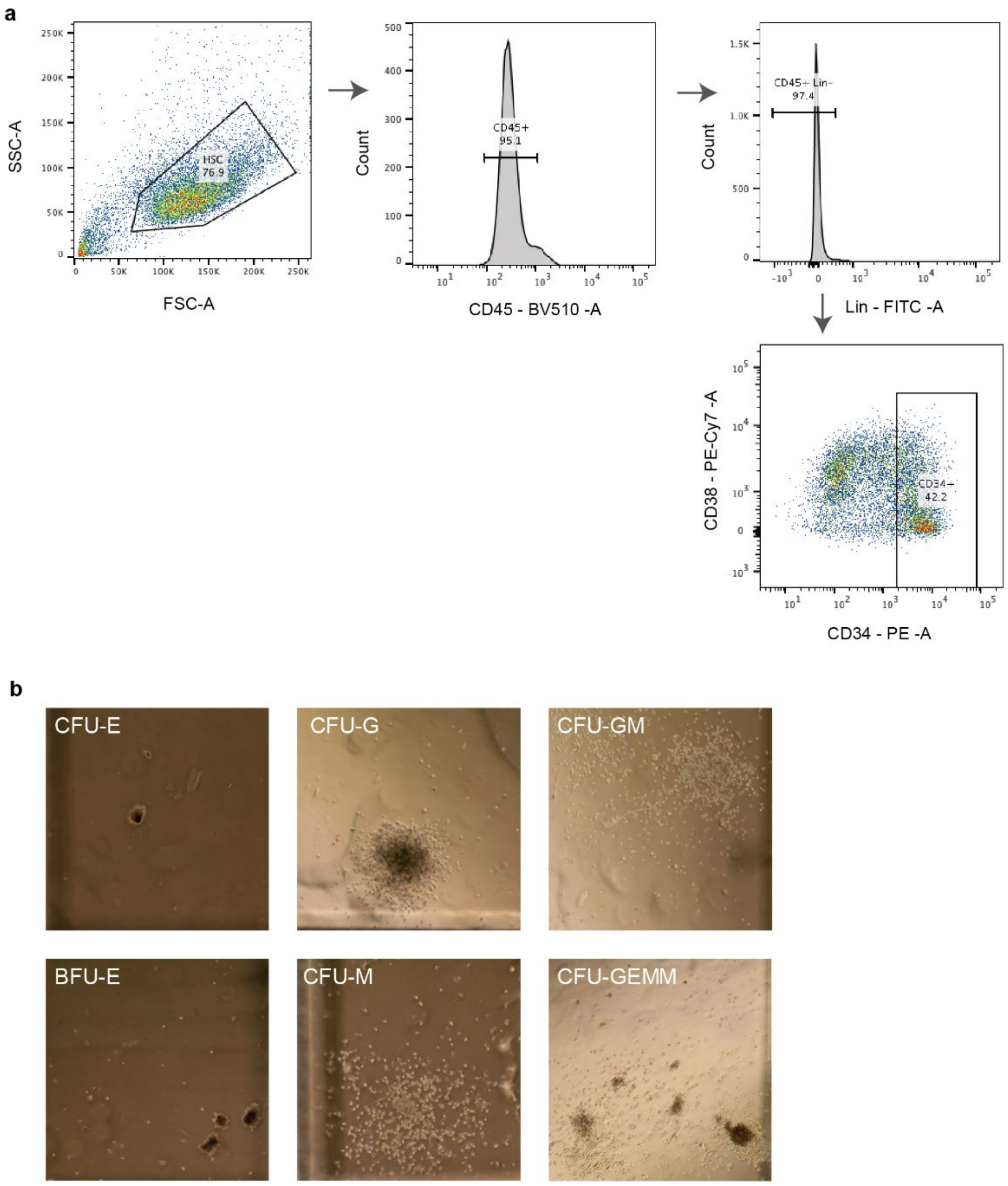


***Supplementary Figure 11 | LTC-IC FACS gating strategy and colony phenotyping. a**, Sample data taken from -gel conditions, patient 1. FSC-A versus SSC-A plot used to identify viable cells; gate added on CD45$^{+ve}$ cells to remove any PerSCs; Lin$^{-ve}$ gate used to remove any committed progenitors and final gate added on CD34$^{+ve}$ population. These cells were collected and used in the LTC-IC assay. **b**, Representative colony types in colony forming unit (CFU) assay after 7 days. CFU-E; contain ~8-200 haemoglobinised erythroblasts. BFU-E; clusters of erythroid progenitors with high proliferative capacity. CFU-G; homogenous population of granulocyte progenitors. CFU-M; homogenous population of macrophage progenitors. CFU-GM; heterogenous population of macrophages and granulocytes. CFU-GEMM; multilineage progenitors that give rise to erythroid, granulocyte, macrophage and megakaryocyte lineages.*

**a**

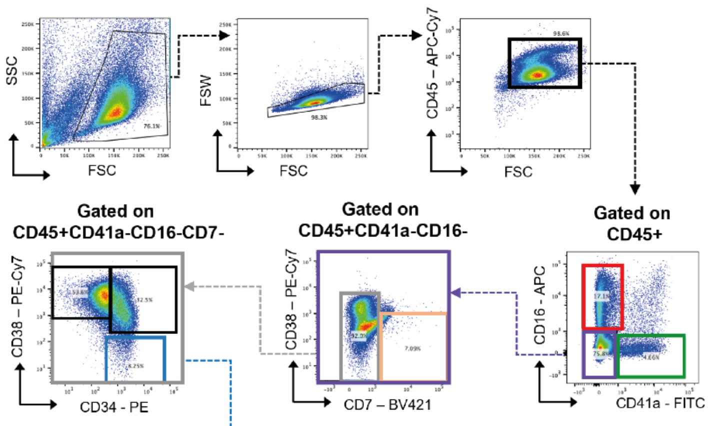


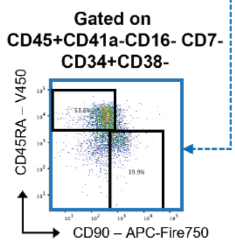


| Gate | Population |
|---|---|
| **CD45+** | **Hematopoietic** |
| **CD45+ CD41a+** | **Megakaryocyte** |
| **CD45+ CD16+** | **Monocytes, B cells, Neutrophils** |
| **CD45+ CD7+** | **T cells** |
| **CD45+CD41a-CD16-CD7-CD38+CD34+** | **Multipotent Progenitors** |
| **CD45+CD41a-CD16-CD7-CD38+CD34-** | **Myeloid Progenitors** |
| **CD45+CD41a-CD16-CD7-CD38-CD34+** | **CD34+ Progenitor Population** |
| **CD45+CD41a-CD16-CD7-CD38-CD34+CD90+CD45RA-** | **LT-HSC** |
| **CD45+CD41a-CD16-CD7-CD38-CD34+CD90-CD45RA+** | **ST-HSC** |

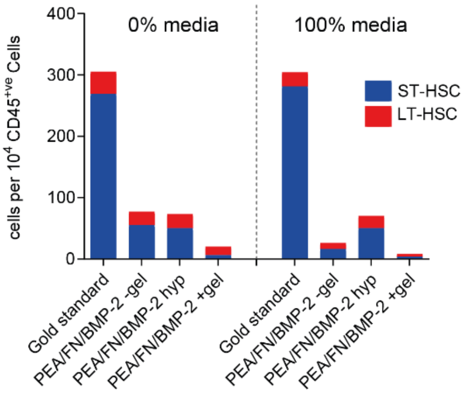


**b**

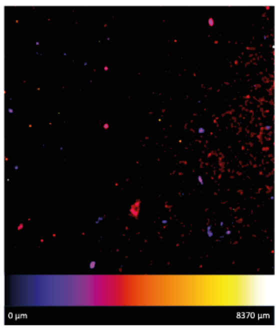


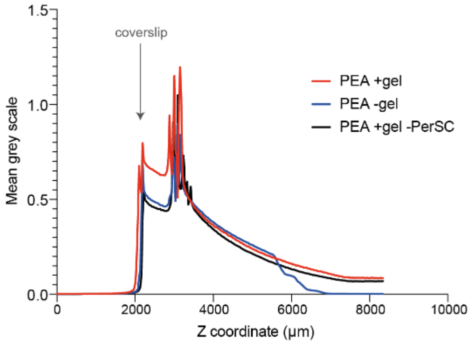


***Supplementary Figure 12 | LT-HSC and ST-HSC phenotyping by flow cytometry. a**, Gating strategy for extended flow cytometry panel comparing markers for LT-HSCs and ST-HSCs. Graph shows number of LT-HSC/ST-HSC per CD45+ cells and demonstrates expansion of the ST-HSC compartment in the gold standard, whereas PEA +gel maintains a population of LT-HSCs. **b**, HSC migration in collagen gels was assessed via NucRed staining of HSCs and distribution in the gels measured via z-projections at day 5. The histogram shows most HSCs migrate down toward the coverslip, regardless of the presence of PerSCs in the niche models.*

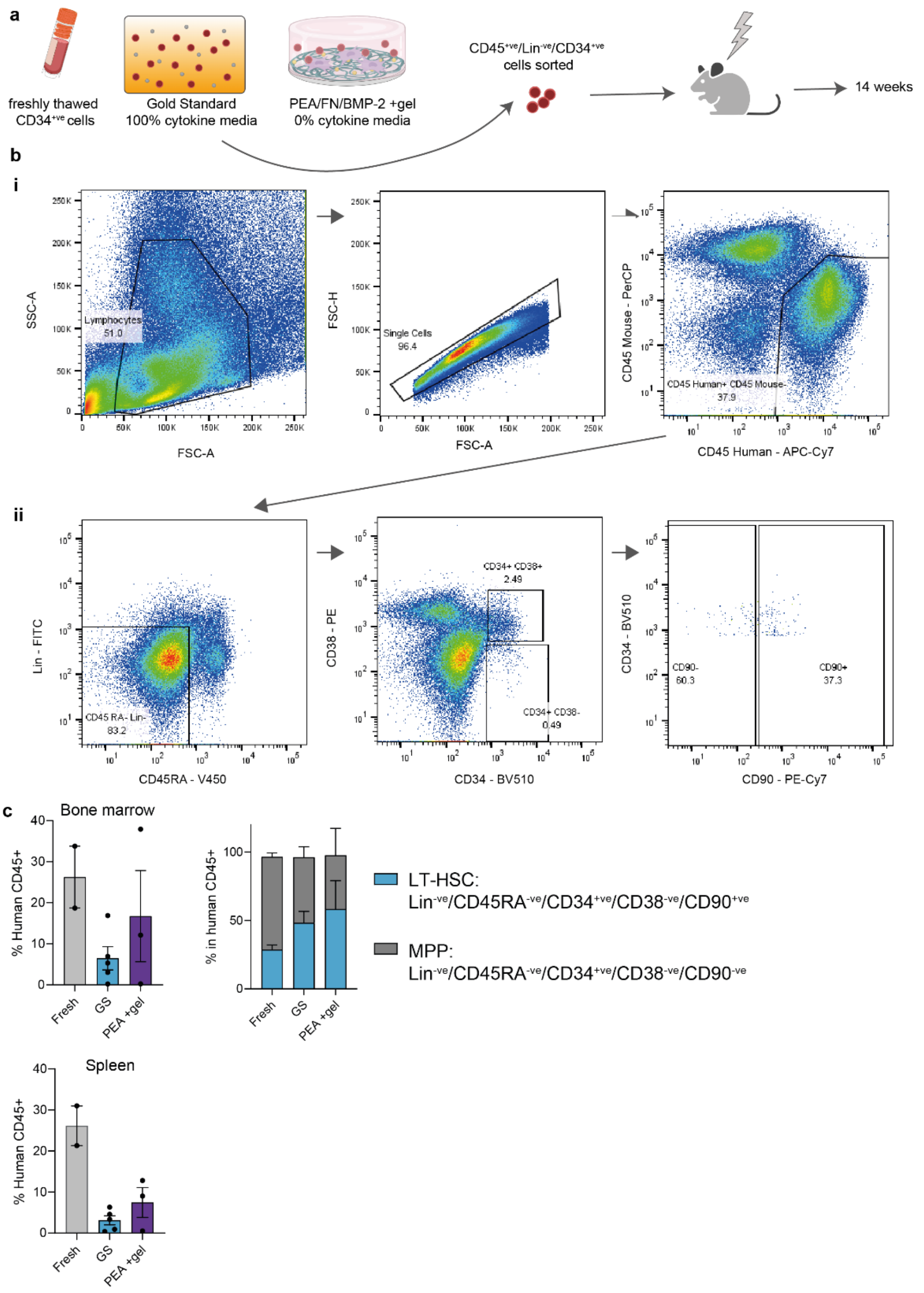


***Supplementary Figure 11 | In vivo reconstitution of CD34$^{+ve}$ cells from bioengineered niches*. *a*, schematic shows experimental set up, briefly CD34$^{+ve}$ cells were seeded in gold standard (GS) media with 100% cytokines, or in PEA/FN/BMP-2 +gel niches in 0% cytokine media. After 5 days CD45$^{+ve}$/Lin$^{-ve}$/CD34$^{+ve}$ cells were sorted and transplanted into NOD-RAG-$\gamma c^{-/-}$ mice, to establish the lineage potential of human HSCs derived from indicated culture conditions in vivo. *b*, Representative gating strategy i. for analysis of % human CD45$^{+ve}$ cells in peripheral blood, and ii. for stem cells marker analysis. *c*, Percentage human CD45$^{+ve}$ cells in the BM and spleen, with comparison of % LT-HSCs and multipotent progenitors (MPPs) within the CD45$^{+ve}$ population of the BM.*

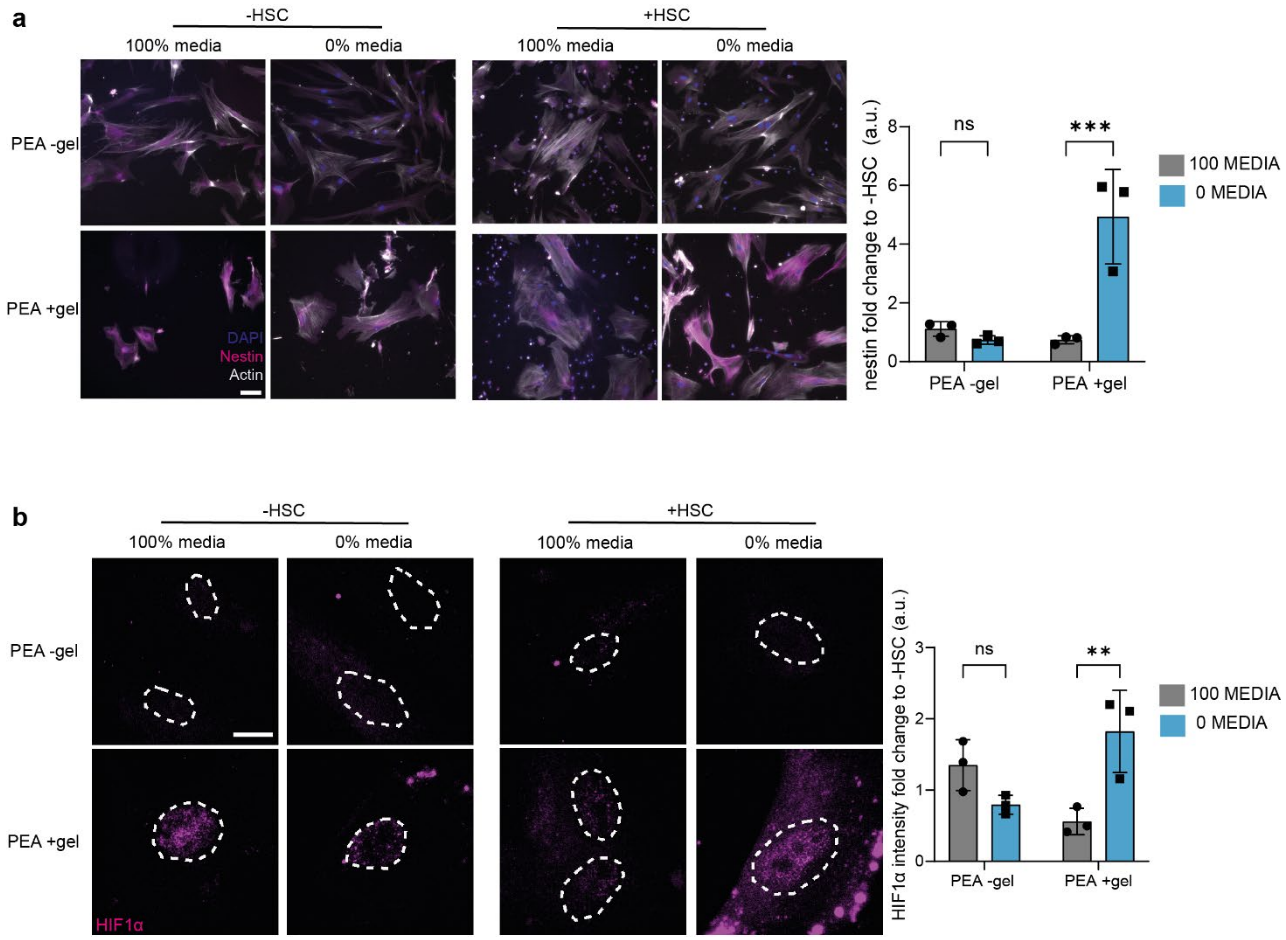


***Supplementary Figure 12 | PerSC phenotype after HSC co-culture at day 19***. ***a***, *Nestin and* ***b***, *HIF1a co-localisation to the nucleus are both significantly increased in PerSCs during HSC co-culture in PEA/FN/BMP-2 +gel (PEA +gel) niches, only when 0% cytokine media is used (the LT-HSC supportive microenvironment).* ***a***, *scale bar = 100 µm, magenta = nestin, grey = actin, blue = DAPI.* ***b***, *scale bar = 100 µm, dashed white line represents nuclear mask detected by DAPI staining, magenta = HIF1a. n = 3 material replicates from 1 biological donor. Actin/DAPI outlines were used to measure individual cell/nuclei nestin/ HIF1a integrated intensity with background correction.*

a

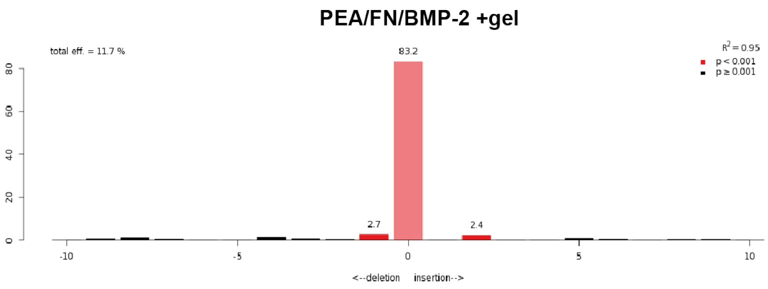


b

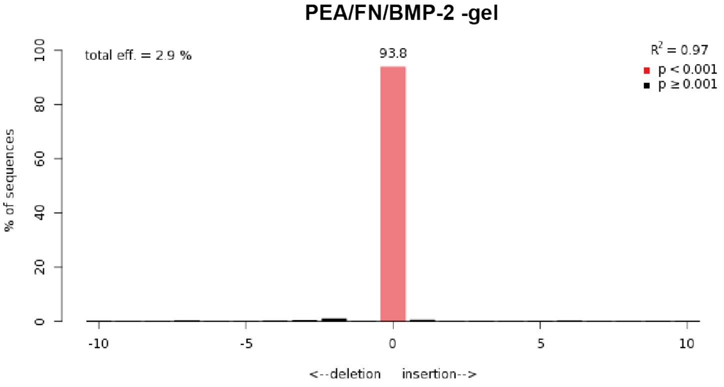


***Supplementary Figure 13 | TIDE analysis of CRISPR-edited HSCs after co-culture in niches.*** *CD34$^{+ve}$ cells were edited using CRISPR targeting the AAVS1 locus and co-cultured in control gold standard media, or in niche models, for 5 days post-editing. Culture in PEA+gel niches enhanced editing,* ***a*** *&* **b***, shows indel spectrum for PEA+gel and PEA-gel HSCs cultured in 0% media, respectively.*

**Supplementary Table 1| Differentiation-related genes of interest.** TGF, transforming growth factor; MSC, mesenchymal stem cell; BMP, bone morphogenetic protein, SMAD, mothers against decapentaplegic; ERK, extracellular related kinase.

| **Gene** | **Definition** | **Summary** | **Supp refs** |
|---|---|---|---|
| *BDNF* | Brain-Derived Neurotrophic Factor | Neurogenesis modulator. | [1,2] |
| *BMP2* | Bone Morphogenetic Protein-2 | TGF-β superfamily member signalling molecules, involved in bone formation. | [3–5] |
| *RUNX2* | Runt-Related Transcription Factor-2 | Transcriptional master regulator of osteogenesis in MSCs. | [6–8] |
| *SMAD4* | SMAD Family Member 4 | Involved in canonical BMP signalling. | [9] |
| *BMP6* | Bone Morphogenetic Protein-6 | TGF-β superfamily member signalling molecules, expressed in hypertrophic cartilage. | [5,10] |
| *GFAP* | Glial Fibrillary Acidic Protein | Intermediate filament protein found in astroglia cytoskeleton. | [11,12] |
| *BGLAP* | Osteocalcin | Expressed and secreted by osteoblasts. | [8,13–15] |
| *BMP4* | Bone Morphogenetic Protein 4 | TGF-β superfamily member, role in embryogenesis and osteogenesis. | [5,16,17] |
| *ALPP* | Alkaline Phosphatase | Metalloenzyme, catalyses hydrolysis of phosphoric acid monoesters. Involved in bone formation. | [7,8] |
| *SP7* | Osterix | Bone-specific transcription factor. Required for osteoblast differentiation. | [18] |
| *ADIPOQ* | Adiponectin | Adipocyte-specific secretory protein. | [19–21] |
| *TGFB3* | Transforming Growth Factor Beta 3 | Bind TGF-β receptors and activate SMAD signalling. | [9,22] |
| *EGF* | Epidermal Growth Factor | Role in growth, proliferation and differentiation of several cell types. | [23] |
| *VWF* | Von Willebrand Factor | Haemostatic plasma glycoprotein, expressed by endothelial cells and some HSCs. | [22,24,25] |
| *SPARC* | Osteonectin | Matrix-associated protein required for collagen calcification in bone. | [26] |
| *BMPR1A* | Bone Morphogenetic Protein Receptor Type 1A | Transmembrane serine/threonine kinase receptors for TGF-β superfamily ligands. Involved in bone formation. | [7,27] |
| *ALPL* | Alkaline Phosphatase, Biomineralization Associated | Metalloenzyme, catalyses hydrolysis of phosphoric acid monoesters. Involved in bone formation. | [7,8] |

| | | | |
|---|---|---|---|
| *PPARG* | Peroxisome Proliferator Activated Receptor Gamma | Master regulator of adipocyte differentiation. | [28] |
| *BMPR2* | Bone Morphogenetic Protein Receptor Type 2 | Transmembrane serine/threonine kinase receptors for TGF-β superfamily ligands. Involved in SMAD signalling. | [29,30] |
| *COL1A1* | Collagen Type I Alpha Chain | Pro-alpha chains of type I collagen. | [7] |
| *SOX9* | SRY-Box Transcription Factor 9 | Master regulator of chondrogenic differentiation in MSCs. | [31] |
| *BMP7* | Bone Morphogenetic Protein 7 | TGF-β superfamily member signalling molecule, promotes osteoblast differentiation. | [32,33] |
| *FGF10* | Fibroblast Growth Factor 10 | Role in cell division, proliferation, differentiation and survival. Important in development. | [34] |
| *BMPR1B* | Bone Morphogenetic Protein Receptor Type 1B | Transmembrane serine/threonine kinase receptors for TGF-β superfamily ligands. Involved in SMAD signalling. | [35] |
| *SPP1* | Osteopontin | Non-collagenous protein present in bone matrix. | [8,36,37] |
| *FGF2* | Fibroblast Growth Factor 2 | Role in cell division, proliferation, differentiation and survival. Can signal through ERK to activate RUNX2. | [38] |

**Supplementary Table 2 | Niche-related genes of interest.** BM, bone marrow; HSC, hematopoietic stem cell; MSC, mesenchymal stem cell; ECM, extracellular matrix; VEGF, vascular endothelial growth factor; ESC, embryonic stem cell.

| **Gene** | **Definition** | **Summary** | **Supp refs** |
|---|---|---|---|
| *CDH2* | Cadherin 2 | Expressed by osteolineage cells in the BM niche. | [39–43] |
| *ALCAM* | Activated Leukocyte Cell Adhesion Molecule | MSC/pericyte marker. | [24,44] |
| *ITGAV* | Integrin Subunit Alpha V | Expressed by nestin+ perivascular stromal cells in the BM niche. | [44] |
| *CSPG4* | Chondroitin Sulphate Proteoglycan 4 (NG2; Nerve/Glial antigen 2) | Expressed by arteriole lining pericytes in the BM niche. | [45–47] |
| *JAG1* | Jagged Canonical Notch Ligand 1 | Expressed by BM stromal cells and osteoblasts. May activate Notch signalling in HSCs. | [47,48] |
| *KITLG* | KIT Ligand | Stem Cell Factor (SCF); HSC maintenance growth factor produced by Lepr+ and Nestin+ stromal and endothelial cells in the niche. | [49] |
| *LIF* | Leukaemia Inhibitory Factor | Interleukin-6 family cytokine, important in haematopoietic differentiation and regulation. | [50,51] |
| *ITGB1* | Integrin Subunit Beta 1 | Expressed by HSCs and MSCs in the BM niche. | [4,52,53] |

| | | | |
|---|---|---|---|
| *PDGFRA* | Platelet Derived Growth Factor Receptor Alpha | Expressed by a subset of CD51+ stromal cells in the BM niche that correspond to increased nestin expression. | [44] |
| *MMP2* | Matrix Metallopeptidase 2 | Shown to release CXCL12 from the ECM. | [54–56] |
| *ICAM1* | Intercellular Adhesion Molecule 1 | Expressed by immunomodulatory MSCs and by sinusoidal endothelial cells in the BM niche. | [22,57] |
| *CXCL12* | C-X-C Motif Chemokine Ligand 12 | Stromal Cell-Derived Factor 1(SDF-1). HSC regulatory cytokine produced by nestin+ perivascular cells in the BM niche. | [46] |
| *ITGAX* | Integrin Subunit Alpha X | Expressed in the BM niche by lymphocytes. | [58] |
| *PDGFRB* | Platelet Derived Growth Factor Receptor Beta | Expressed by endothelial cells lining BM arterioles and sinusoids. | [49] |
| *LEPR* | Leptin Receptor | Expressed by sinusoid-associated cells in the bone marrow niche. | [45–47] |
| *VCAM1* | Vascular Cell Adhesion Molecule 1 | Involved in blood cell retention in the BM niche, ligands is integrinβ1. | [46,49] |
| *NOTCH1* | Notch Receptor 1 | Roles in osteoblast and hematopoietic cell differentiation and maturation. | [59–61] |
| *MCAM* | Melanoma Cell Adhesion Molecule (CD146) | Marker for pericytes and stromal stem cells. | [24,44] |
| *NCAM1* | Neural Cell Adhesion Molecule 1 | Considered an early neural progenitor marker – correlated with nestin expression. | [62] |
| *NES* | Nestin | Nestin+ pericytes and stromal cells regulate HSC activity in the BM niche. | [44,45,47,63–65] |
| *THY1* | Thy-1 Cell Surface Antigen | CD90; Involved in early BM niche development, used as MSC/HSC marker. | [66–68] |
| *CSF2* | Colony Stimulating Factor 2 | Granulocyte macrophage-colony stimulating factor (GM-CSF). Produced in multiple cell types the BM niche. | [69,70] |
| *KDR* | Kinase Insert Domain Receptor | VEGF receptor. MSC marker. | [71,72] |
| *THPO* | Thrombopoietin | Osteoblast-derived HSC maintenance cytokine. | [73] |
| *NGFR* | Nerve Growth Factor Receptor | MSC marker. | [74] |
| *VIM* | Vimentin | Intermediate filament protein expressed in mesenchymal cells. Polymerisation partner of nestin. | [75] |
| *POU5F1* | Octamer-Binding Protein 4 | Regulates self-renewal and differentiation in MSCs. Shown to be regulated by hypoxia in cultured ESCs. | [76,77] |

| | | | |
|---|---|---|---|
| *VEGFA* | Vascular Endothelial Growth Factor A | Regulator of blood vessel formation. Required for HSC survival. Produced in hypoxia. | 78 |

***Supplementary Table 3| Flow cytometry antibodies for PerSC phenotyping.***

| Marker | Fluorophore | Clone | Supplier |
|---|---|---|---|
| Lepr | APC (panel 1) | REA361 | Miltenyi Biotech |
| CD51 | APC (panel 2) | NKI-M9 | Biolegend |
| CD90 | APC-Cy7 (panel 1) | 5E10 | ThermoFisher, eBioscience |
| CD31 | APC-Cy7 (panel 2) | WM59 | ThermoFisher, eBioscience |
| CD29 | FITC (panel 1) | TS2/16 | ThermoFisher, eBioscience |
| NG2 | PE (panel 1) | 7.1 | ThermoFisher, eBioscience |
| CD140a | PE (panel 2) | 16A1 | ThermoFisher, eBioscience |
| CD146 | PerCP-Cy5.5 (panel 1) | P1H12 | Biolegend |
| CD166 | PerCP-eFluor710 (panel 2) | 3A6 | ThermoFisher, eBioscience |
| CD140b | PE-Cy7 (panel 2) | APB5 | ThermoFisher, eBioscience |
| CD105 | eFluor 450 (panel 2) | SN6 | ThermoFisher, eBioscience |

***Supplementary Table 4| Flow cytometry antibodies for HSC phenotyping, LTC-IC and in vivo assay sorting.***

| Marker | Fluorophore | Clone | Supplier |
|---|---|---|---|
| CD34 | PE | 4H11 | ThermoFisher, eBioscience |
| CD45 | APC-Cy7 | 2D1 | BD Biosciences |
| CD38 | PE-Cy7 | HB7 | ThermoFisher, eBioscience |
| Lineage cocktail | FITC | - | ThermoFisher, eBioscience |
| CD16 | APC | 3G8 | ThermoFisher, eBioscience |
| CD7 | BV421 | 562635 | ThermoFisher, eBioscience |
| CD90 | APC-Fire750 | 5E10 | Biolegend |
| CD45RA | V450 | HI100 | ThermoFisher, eBioscience |
| CD41a | FITC | HIP8 | ThermoFisher, eBioscience |
| CD45 | BV510 | 2D1 | Biolegend |
| CD90 | PerCP-Cy5.5 | 5E10 | ThermoFisher, eBioscience |
| CD45RA | APC-Cy7 | HI100 | ThermoFisher, eBioscience |

*Supplementary Table 5 | Flow cytometry antibodies for HSC phenotyping for in vivo experiments.*

| **Marker** | **Fluorophore** | **Clone** | **Supplier** |
|---|---|---|---|
| Lin | FITC | - | BD Biosciences |
| CD34 | BV510 | 581 | Biolegend |
| CD45 (human) | APC-CY7 | 2D1 | BD Biosciences |
| CD45 (Mouse) | PerCP Cy5.5 | 30-F11 | BD Biosciences |
| CD38 | PE | HIT2 | BD Biosciences |
| CD90 | PE-Cy7 | 5E10 | BD Biosciences |
| CD123 | PE Cy7 | 7G3 | BD Biosciences |
| CD11b | Pacific blue | ICRF44 | BD Biosciences |
| CD8 | BV510 | SK1 | BD Biosciences |
| CD3 | PE | 17A2 | BD Biosciences |
| CD45RA | Efluor 450 | HI100 | Thermofisher |
| CD56 | PE-Cy7 | B159 | BD Biosciences |
| CD19 | APC | HIB19 | BD Biosciences |
| FC Block (CD16/32) | - | 2.4G2 | BD Biosciences |
| UltraComp eBeads | - | - | Themofisher |

### Supplementary methods

#### Trilineage differentiation of PerSCs

PerSCs at ≤passage 5 were cultured in differentiation medias for 4 weeks: osteogenic media - DMEM with 10% FBS, 0.1 µM dexamethasone (Sigma, D2915), 100 µM Ascorbate-2-phosphate (Sigma, A8960), 10 mM β-Glycerophosphate disodium salt hydrate (Sigma, G9422); Adipogenic media – DMEM with 10% FBS, 1 µM dexamethasone, 500 µM 3-isobutyl-1-methylxanthine (IBMX) (Sigma, I5879), 1.8 µM Insulin (Sigma, I9278), 100 µM indomethacin (Sigma, I7378); Chondrogenic media – 100 nM dexamethasone, 100 nM Ascorbate-2-phosphate, 1% (v/v) insulin, transferrin, selenium (Thermo Fischer, 41400-045), 40 mg/mL L-proline, 10 ng/mL TGFβ3 (Peprotech, 100-36E). Cells were then fixed using 10% formaldehyde for 15 mins at RT. After washing with PBS, osteogenic fixed cells were stained with 2% (w/v) Alizarin Red solution (pH 4.1 to pH 4.3) for 15 min at room temperature. After staining, cells were washed in deionized water. For adipogenic staining, fixed cells were washed with distilled water three times, rinsed with 60% (v/v) isopropanol. Oil Red O solution was then added to the cells, and cells were incubated at room temperature for 15 min. Dye solution was removed, and cells were washed again with 60% (v/v) isopropanol, washed three times in distilled water. For chondrogenic staining, cells were fixed with 0.1 % glutaraldehyde in PBS for 20 min at RT. Rinsed 3 x PBS, rinsed 1 x with 1 % acetic acid solution for 10–15 s and stained with 0.1 % Safranin O solution (Merck) for 5 min. Cells were then washed 3 x PBS. Cells were then imaged on an inverted microscope (Olympus, PA, USA) operated through Surveyor software (v.9.0.1.4, Objective Imaging, Cambridge, UK). Images were processed using ImageJ [v.1.50g, National Institutes of Health (NIH), USA].

#### Lactate assay

PerSCs were cultured in niche systems in DMEM with 2% dialyzed FBS (ThermoFisher, A3382001), cell media supernatant was aspirated at 7 and 14 days and stored at -20°C. Lactate levels in supernatant were then measured using the Lactate-Glo™ assay (Promega, J5021) as per manufacturers protocol. Briefly, samples were diluted 1:100 and mixed 1:1 with Lactate Detection Reagent in 96-well plate and incubated for 1 h at RT. Luminescence was read using PHERAstar FSX microplate reader (BMG Labtech).

#### In-cell western assay for LDH

PerSCs were cultured in niches for 7 days then fixed using 10% formaldehyde for 15 min. The cells were then permeabilized with 0.5% Triton-X for 5 mins and blocked using 0.5% milk protein in 1X PBS. Anti-lactate dehydrogenase (LDH) [EP1566Y] (rabbit monoclonal, Abcam, ab52488) was added at 1:200 in 0.5% milk PBS and incubated at 4°C overnight, followed by 5 washes in PBST. CellTag 700 stain (LI-COR, 926-41090) was used as the reference control, and was added with LI-COR anti-rabbit secondary antibody (926-32211), at 1:2000 in 0.5%

milk PBST and incubated at RT on a shaker for 2 h, followed by 5 x 5 min washes with PBST. The quantitative spectroscopic analysis was the carried out using the LI-COR Odyssey Sa.
